# Evidence against the detectability of a hippocampal place code using functional magnetic resonance imaging

**DOI:** 10.1101/229781

**Authors:** Christopher R. Nolan, J.M.G. Vromen, Allen Cheung, Oliver Baumann

## Abstract

Individual hippocampal neurons selectively increase their firing rates in specific spatial locations. As a population these neurons provide a decodable representation of space that is robust against changes to sensory- and path-related cues. This neural code is sparse and distributed, theoretically rendering it undetectable with population recording methods such as functional magnetic resonance imaging (fMRI). Existing studies nonetheless report decoding spatial codes in the human hippocampus using such techniques. Here we present results from a virtual navigation experiment in humans in which we eliminated visual- and path-related confounds and statistical shortcomings present in existing studies, ensuring that any positive decoding results would be only spatial in nature and would represent a true voxel-place code. Consistent with theoretical arguments derived from electrophysiological data and contrary to existing fMRI studies, our results show that although participants were fully oriented during the navigation task, there was no statistical evidence for a place code.

## Introduction

Acquisition of declarative memories is dependent on the hippocampus. Place cells — hippocampal principal cells that exhibit allocentric spatial tuning — provide a clear behavioural correlate with which to interrogate the neuronal dynamics of this region (O’Keefe and Dostrovsky, 1971). Initially discovered in rodents, the existence of place cells has since been confirmed in other species, including humans (Ekstrom et al., 2003). The activity across populations of such cells, as measured with single cell recordings, can be decoded to provide an accurate estimate of an animal’s current position (Brown et al., 1998), and the activity appears to reflect a cognitive map, resilient against changes in any particular internal or external cue. However, the sparse firing and random distribution of spatial tuning amongst the place cell population suggests that any such place code should be impenetrable to current mass imaging technology such as fMRI.

A true place code should be demonstrably selective for position in a mnemonic representation of space rather than particular external or non-mnemonic internal cues such as unique visual patterns or egocentric movement. We are aware of four studies that claim to provide evidence for a voxel place code (Hassabis et al., 2009; Kim et al., 2017; Rodriguez, 2010; Sulpizio et al., 2014). Each experiment involved distinguishing between fMRI scans taken at two or more locations in a virtual arena. All four experiments failed to remove significant visual confounds, either in the form of salient visual landmarks during navigation to a target (Hassabis et al., 2009; Kim et al., 2017; Rodriguez, 2010) or at the target (Rodriguez, 2010; Sulpizio et al., 2014), or as visual panoramas unique to each target location (Kim et al., 2017). We later discuss how these confounds, amongst others, are manifest in each experiment (see Discussion), but note here that any legitimate voxel codes in these experiments could be sensory-driven rather than true place codes.

Beyond experimental design issues, detecting a voxel place code necessitates distinguishing between complex multivariate voxel patterns. Each of the existing four studies uses multivariate pattern analysis (MVPA) techniques to classify voxel patterns as characteristic of particular virtual locations. While these analytics are valid in principle, subsequent statistical inferences cannot necessarily rely on classical assumptions. For example, the practice of submitting within-subject classification results to a second-level t-test to infer group statistics — a technique used by two of the referenced hippocampal studies — has been demonstrated as invalid on information-like measures such as classification results (Allefeld et al., 2016). In particular, the true value of information-like measures cannot be below chance, thereby restricting the null hypothesis to be the total absence of information. Hence even when the null is rejected, the strongest conclusion possible is that there are people in whom information is found, not that the information is prevalent or generalizable (Allefeld et al., 2016). Additionally, such measures violate assumptions of Gaussian or other symmetric null distributions (Stelzer et al., 2013; Brodersen et al., 2013). We found that in three of the existing studies (Hassabis et al., 2009; Rodriguez, 2010; Sulpizio et al., 2014), statistical issues marred the interpretation of any evidence (see Methods and Results).

These concerns motivated us to revisit the question of whether a voxel place code is truly detectable with human fMRI. We had a group of healthy participants perform a virtual navigation task, while undergoing high-resolution3T fMRI. The environment was a circular arena containing two unmarked target locations (see Figure 1a). On each trial, participants were initially shown an orienting landmark and then had to track their position while being passively moved along a curvilinear path to one of the two target locations. During navigation, the participants had to rely solely on their mental representation of the environment and track their position using visual self-motion cues. After arriving at one of the target locations, we probed the participants’ positional knowledge. We then used linear and non-linear multivoxel classification methods to test whether we could distinguish hippocampal fMRI signals corresponding to periods at which participants were present at each of the two target locations.

**Figure 1.**
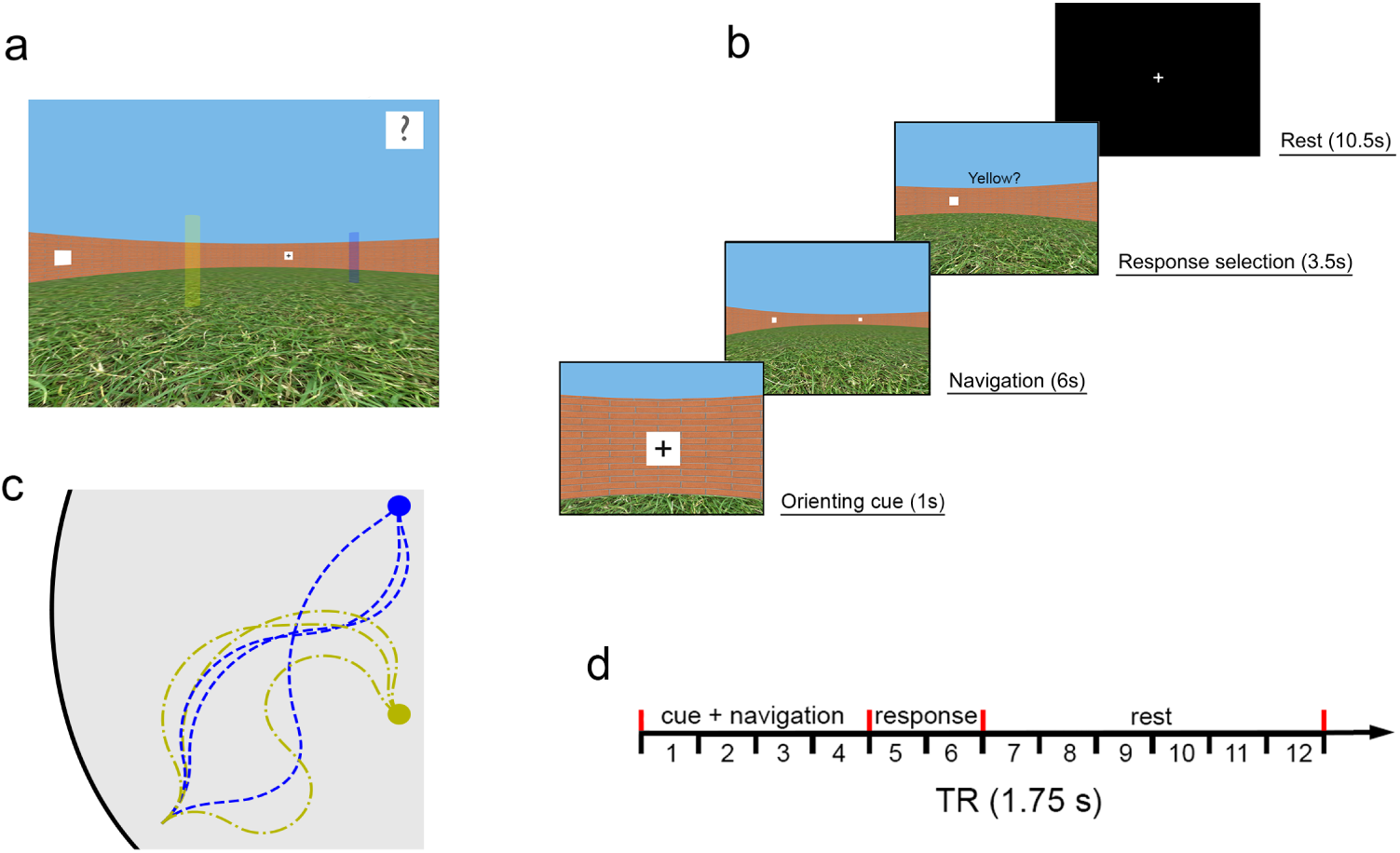
Schematics of the virtual environment and task. **a.** First person view of the environment during the training stage (beacons marking target locations are not visible in the main experiment). **b.** Sequence of events in a typical experimental trial. **c.** Schematics of the path structures used in the experiment. Participants were led to the target location via in total 24 (three paths from each landmark to each beacon) different curvilinear paths of equal length. **d.** Experimental time course of each trial relative to the image acquisition sequence (1.75 s per volume).

## Materials and methods

### Participants

Twenty-one healthy, adult volunteers gave their informed consent to participate in the study, which was approved by the Human Research Ethics Committee of The University of Queensland. The first two participants were only used for pilot testing, to optimise acquisition parameters. One participant was omitted from the data analysis because the behavioural performance was below our a priori criterion of 90% correct responses. The remaining 18 participants (9 females) ranged in age from 18 to 29 years (mean, 21 years) and all were right-handed. Classical sample-size estimation techniques are not applicable to the classification analyses in the present study, however we deemed our sample size sufficient given that three of the four existing studies reported a positive place code effect with fewer subjects (Hassabis et al., 2009; Rodriguez, 2010; Sulpizio et al., 2014).

### Stimuli and procedure

The virtual environment was a circular arena surrounded by a brick wall, with a grass-textured floor and featureless blue sky. The arena wall was 3.0 m high and its diameter was 30.4 m, relative to a 1.7 m observer. Along the wall, four landmarks (white 1.0 m ×1.0 m squares with black symbols: ‘+’, ‘%’, ‘?’, and ‘#’) were located equidistantly (45°, 135°, 225° and 315°). The two beacons (yellow and blue, see Figure 1a) were 3 m tall and 0.5 m in diameter, located at 0° and 180°, and 5 m from the centre of the arena (i.e. 10 m apart from each other).

The task required participants to track their location, while being passively moved (4.2 m/s linear speed) in the absence of orienting landmarks through the environment, therefore relying only on a combination of visual self-motion cues and their mental representation of the landmarks’ locations (see Figure 1b for details of the task sequence). At the beginning of each trial, participants closely faced one of the four peripheral landmarks on the arena wall for one second. Subsequently, all four landmarks were made invisible (i.e. replaced by white placeholders) and participants were turned around and moved for 6 s along a curvilinear path to one of the two unmarked target locations. Participants were led to the target location via 24 different curvilinear paths of equal length (see Figure 1c), so that participants could not infer the target location simply based on the initial landmark cue and the length of the path. After arriving at the target location, participants were prompted to indicate their location within 3.5 s, via a yes/no button response to either the question “Yellow?” or the question “Blue?”, chosen at random. This procedure ensured that the button response was orthogonal to the target location. The response period was followed by a 10.5 s rest period, in which only a white fixation cross on a black screen was shown (see Figure 1b). There were in total 120 trials (60 per target location) split up into five imaging runs, lasting ~8.5 minutes each.

We used the Blender open-source three-dimensional content creation suite (The Blender Foundation)to create the virtual maze and to administer the task. Stimuli were presented on a PC connected to a liquid crystal display projector (1280 × 980 resolution) that back projected stimuli onto a screen located at the head end ofthe scanner bed. Participants laid on their back within the bore of the magnet and viewed the stimuli via a mirror that reflected the images displayed on the screen. The distance to the screen was 90 cm (12 cm from eyes to mirror) and the visible part ofthe screen encompassed ~22.0° × 16.4° of visual angle (35.5 × 26 cm).

Before conducting fMRI imaging, participants were assessed and trained using a three-stage procedure to ensure an adequate level of task performance, which depends on familiarity with the arena layout. These behavioural training sessions were scheduled one to two days before the fMRI scanning session. In the first training stage, participants were allowed to freely navigate the virtual environment for three minutes, using a joystick held in their right hand. During this stage, all four wall landmarks and the two beacons that marked the target locations (yellow and blue) were visible. In the second stage ofthe training only the two beacons and one ofthe peripheral landmarks were visible at a time, and the participants’task was to navigate to the location of one ofthe otherthree landmarks, indicated by a small cue (an image ofthe landmark) atthetopof the computer screen. Each participant completed at least 24 trials of this task. Thethird stage ofthetraining procedure was almost identical to the actual task described earlier, except the yellow and blue beacons marking the two target locations were visible during the first six trials, feedback was provided for 1.5 s after each button press (i.e. “correct”/“incorrect”), and the interval between trials was just 2 s. Each participant completed at least 24 trials of this task. When participants achieved a performance level of >90% correct in the last stage of the training they were admitted to the fMRI session. At the beginning ofthe scanning session, during the acquisition ofthe structural images, participants performed another iteration ofthe trainingtasks to refamiliarize them with the environment.

### MRI acquisition

Brain images were acquired on a 3T MR scanner (Trio; Siemens) fitted with a 32-channel head coil. For the functional data, 25 axial slices (voxel size 1.5 ×1.5 ×1.5 mm, 10% distance factor) were acquired using a gradient echo echoplanar T2*-sensitive sequence (repetition time, 1.75 s; echo time, 30.2 ms; flip angle, 73°; Acceleration factor (GRAPPA), 2; matrix, 128 ×128; field of view, 190 ×190 mm). In each of five runs, 294 volumes were acquired for each participant; the first four images were discarded to allow for T1 equilibration. We also acquired a T1-weighted structural MPRAGE scan. To minimize head movement, all participants were stabilized with tightly packed foam padding surrounding the head.

## Data analysis

### Preprocessing

Image preprocessing was carried out using SPM12 (Wellcome Department of Imaging Neuroscience, University College London). Functional data volumes were slice-time corrected and realigned to the first volume. A T2^*^-weighted mean image of the unsmoothed images was coregistered with the corresponding anatomical T1-weighted image from the same individual. The individual T1 image was used to derive the transformation parameters for the stereotaxic space using the SPM12 template (Montreal Neurological Institute template), which was then applied to the individual coregistered EPI images. Further, to exclude voxels with spurious signals, we removed all voxels with a raw intensity of zero at any time during the timeseries (RH: 0.37 ±0.20%, LH: 0.83 ± 0.54%, RPH: 2.0 ± 1.3%, LPH: 4.4 ± 3.1%; mean ± SD%, n = 18).

Two alternative approaches of detrending were used to assess their potential differential effect on decoding performance. (1) To make use of global information about unwanted signals, images were detrended using a voxel-level linear model of the global signal (LMGS; Macey et al. (2004)) to remove high-frequency as well as low-frequency noise components due to scanner drift, respiration, or other possible background signals. (2) To remove spatiotemporally confined signal drift and artefacts, runwise polynomial detrending was performed on region of interest (ROI) data (see below). By default, second order polynomial detrending was used (SPM, Wellcome Department of Imaging Neuroscience, University College London, London, UK).

Based on existing evidence that in humans the right hippocampus should be the most likely region to produce a place code (Burgess et al., 2002), we used the AAL atlas (Tzourio-Mazoyer et al., 2002) and WFU pickatlas tool (Maldjian et al., 2003) to generate a right hippocampal (RH) ROI mask. For additional control analyses, we also generated ROI masks for the left hippocampus (LH), left parahippocampal gyrus (LPH), and right parahippocampal gyrus (RPH). The masks were separately applied to the 4D timeseries using Matlab 2015b (Mathworks, Inc.).

### Multivariate pattern classification

We performed a ROI-based multivariate analysis (Haynes, 2015) designed to test whether fMRI activation patterns in the human hippocampus carry information about the participants’ position in the virtual environment. The fMRI BOLD signal has an inherent delay relative to stimulus onset of ~2 s until it increases above baseline, and ~5 s to peak response (Huettel et al., 2014). To account for this delay, we selected for the analysis the volumes corresponding to the period of 3.5–5.25 s after participants arrived at the target location (i.e. fMRI TR #7 of our 12-TR trial structure, see Figure 1d). The volume selection approach is analogous to that employed by Hassabis et al. (2009) and Rodriguez (2010).

The goal of our multivariate analysis was to test whether we could classify the virtual location of the participant using the selected volumes. The classification was performed using a linear support vector machine (Haynes, 2015), denoted here as LSVM, implemented in Matlab 2015b (Mathworks Inc.). Two data sets were constructed, one with correct labels (location 1 or location 2), and one with randomly shuffled labels. Each data set was then randomly partitioned into 10 subgroups (or folds), split evenly between its class labels (stratification). The classifier was trained on 9 folds (training data), and its performance cross-validated on the remaining fold (withheld test data), once for each of the 10 possible combinations of train and test folds. We repeated this procedure 1,000 times for each participant (i.e. 1,000 random 10-fold stratified cross validations), which allowed us to estimate the distribution of classification accuracy with (true class labels) and without (shuffled class labels) class information, as well as the distribution of classification accuracy associated with randomly partitioning the data, referred to here as partition noise. Estimating a distribution for partition noise is an additional step from standard application of SVM to MVPA, where typically a single partition of the correct label data is used. A major goal of MVPA is to determine whether novel multivoxel patterns can be used to predict their true class labels, and there is no way to know a priori how any particular choice of trial assignment among folds affects such predictive capability. Our 1,000 random partitions of the data using true class information allows us to characterise this partition noise distribution.

### Positive control and additional verification analyses

As a direct comparison using the same data and preprocessing steps, we replicated the ROI-based SVM analysis to classify two distinct phases within each trial, which we expected to be different at the voxel level (i.e. a positive control). Given that the right hippocampus is known to show task-related activity during spatial navigation tasks (Baumann et al., 2010, 2012; Baumann and Mattingley, 2013), we hypothesized that the hippocampus should express differential fMRI activity patterns during the navigation period of our task compared to the rest period. Taking the delay in the BOLD response into account, we chose fMRI image #4 (navigation) and #12 (rest) of our 12-image trial structure for this comparison (see Figure 1d).

In addition, to eliminate the possibility that negative results could be due to our choice of preprocessing methods, classifier, brain region or fMRI images (i.e. time to signal peak) we conducted several additional analyses to verify the null results. First, to exclude that a particular choice of signal detrending was suboptimal, we performed the same analysis using both LGMS and 2nd order polynomial detrending (see *Preprocessing*). Second, to exclude the possibility that image smoothing may have impaired the discriminability of the fMRI signal we repeated the analysis using unsmoothed images (Kamitani and Sawahata, 2010). Third, we explored whether there was any decodable signal in the left hippocampus (LH ROI). Fourth, to test whether decoding of location information could be improved by averaging fMRI signals over a longer period (i.e. several images), we conducted analyses averaging two (i.e. image #7–#8), as well as three consecutive fMRI images (i.e. image #7–#9). In total, this yielded 24 classification analyses. Finally, to investigate whether there could be voxel place codes that are non-linearly separable, we repeated the same analyses using a radial basis function (Gaussian) SVM (Song et al., 2011), denoted here as RSVM.

### Multivariate searchlight analysis

In addition to the ROI-based classification approach, we also employed so-called searchlight decoding (Kriegeskorte et al., 2006). In this approach, a classifier is applied to a small, typically spherical, cluster of voxels (i.e. the so-called searchlight). The searchlight is then moved to adjacent locations and the classification repeated. This approach has the advantage that the dimensionality of the feature set is reduced, i.e. the multivariate pattern consist of fewer voxels, and makes the analysis more sensitive to information contained in small local volumes. We followed the searchlight and detrending methods of Hassabis et al. (2009), using spherical searchlights of 3-voxel radius (comprising a maximum of 121 voxels), on run-wise linearly detrended data. LSVM was applied, using 100 random 10-fold stratified cross validations for each searchlight, both with and without class label information. Each label shuffle was identical amongst all searchlights to be compatible with subsequent population inferencing and correction for multiple comparisons (Allefeld et al., 2016).

We further included left and right parahippocampal regions in the searchlight analysis in order to compute differences in proportions of searchlights exceeding a classification accuracy threshold following Hassabis et al. (2009). This analysis quantifies the difference between the proportion of searchlights in the hippocampal and parahippocampal regions which exceeded the 95^th^ percentile classification threshold computed from shuffled location labels. To determine if the difference in proportions was greater than expected by chance, Hassabis et al. (2009) estimated the standard error of the difference-of-proportions using a standard result, implicitly assuming statistical independence between searchlight accuracies (but see *Evaluation of analysis used in Hassabis et al. (2009)* for further details on the problems of this assumption). Due to the computing load, this analysis was implemented in Python v3.5 on a 300-node cluster.

### Population inference using a permutation-based approach

For population inference, we followed the nonparametric, permutation-based approach of Allefeld et al. (2016). Allefeld and colleagues provided strong arguments that the random-effects analysis implemented by the commonly used t-test fails to provide population inference in the case of classification accuracy or other‘information-like’ measures, because the true value of such measures can never be below chance level, rendering it effectively a fixed-effects analysis. The reason is that the mean classification accuracy will be above chance as soon as there is an above-chance effect in only one person in the sample. As a result, t-tests on accuracies will with high probability yield ‘significant’ results even though only a small minority of participants in the population shows above-chance classification.

A further advantage of the approach of Allefeld et al. (2016) is the ability to estimate the population prevalence when the prevalence null hypothesis is rejected. This enables direct quantification of the generalisability of a positive finding in the population.

Briefly, first level permutations (within-participant) were classification accuracies where class labels are randomly shuffled, together with one classification accuracy with correct labels. Second level permutations (between-participant) were random combinations of first level permutations across participants, with one ofthe second level permutations consisting of accuracies from all correct labels (to avoid p-values of zero). The minimum statistic was used across subjects for each comparison (e.g. searchlight or ROI), and for each second-level permutation. For each second-level permutation, the maximum statistic across comparisons was computed to correct for multiple comparisons (Allefeld et al., 2016; Nichols and Holmes, 2001). Since the maximum statistic does not depend on the amount or nature of statistical dependence between comparisons, it is applicable to classification accuracies of overlapping regions such as searchlights (Allefeld et al., 2016; Nichols and Holmes, 2001). By the same reasoning, it is also applicable to multiple comparisons across different analyses ofthe same ROI, such as SVM classification following different preprocessing methods. Here, we computed the maximum statistic across all ROIs and preprocessing methods *(Extended analysis of negative results)*, and also the maximum statistic across searchlights in each ROI *(Multivariate searchlight analysis*).

### Stochastic binomial model for shuffled labels

We developed a stochastic binomial model of classification accuracy based on the null hypothesis, and crossvalidation analysis parameters. Each test volume was assumed to be classified stochastically with classification success governed only by the null hypothesis probability *p_0_*. For *k*-fold cross-validation (*k*-fold CV), there are *n_f_* = *N_T_*/*k* binary choices for each of *k* folds, averaged to give the accuracy of a single partition set (stratified, non-overlapping hold-out sets). Assuming the training data is entirely devoid of information, then performance on test data must be at chance, i.e.,

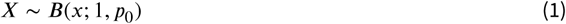

The sample probability of a successful prediction per fold is the number of successful predictions averaged over each fold, i.e.,

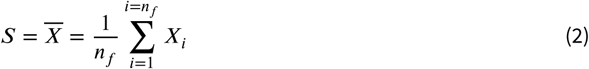

Then the variance ofthe prediction success per trial is

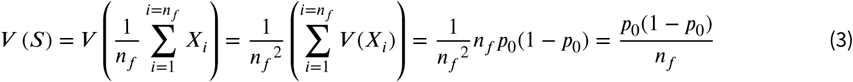

assuming statistical independence between scores within a fold. For truly random partitions and large *N_T_*, this seems a good approximation since volumes in close temporal proximity are rare. Thus if the training data is not informative, then the test data are all essentially independent.

The SVM’s *k*-fold CV accuracy from each random partition is the prediction success averaged over all *k* folds. It is tempting to estimate the variance ofthe average prediction success as

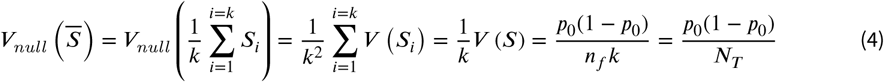

by assuming that folds are statistically independent. The problem is that although folds are predicted based on uninformative training data, uninformative is *not* the same as independent. This is because two training sets overlap by (*N_T_* − 2*n_f_*)/(*N_T_* –*n_f_*) since the data points are drawn from the same set.

The more general form of Equation 4 accounts for covariance terms, i.e.,

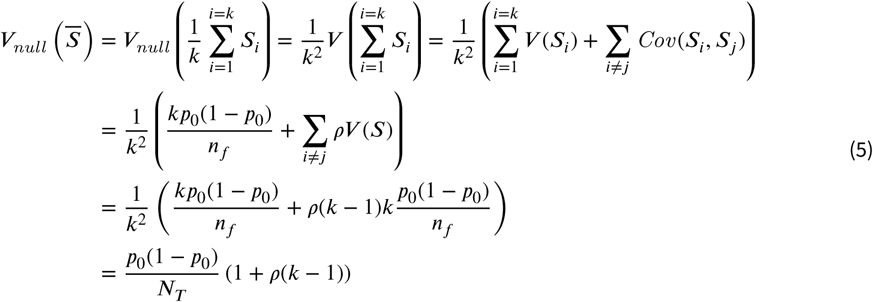

where the correlation coefficient

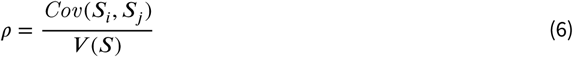

remembering that *V*(*S_i_*) = *V*(*S_j_*) = *V*(*S*). Thus the variance of the null distribution can be written as a function of the null hypothesis probability *p_0_* and the CV parameters, i.e.,

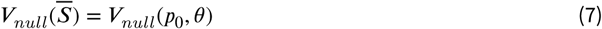

where the CV parameter *Θ* = (*N_T_*, *k*). At present, the correlation coefficient is found empirically assuming each voxel’s signal is independent, normally distributed random noise. Using synthetic noise data instead of fMRI data guarantees there is no classifiable signal in keeping with the null hypothesis, and also enables predictions to be made when designing new experiments. We generated 10^5^ noise data sets, *n_vox_* = 3053 (for RH), *N_T_* = 120, *k* = 10. Using LSVM, *ρ* = 0.0741.

For computational efficiency, we used a Gaussian approximation of the binomial model

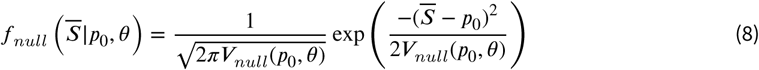

### Stochastic binomial model for true labels

To model the partition noise of individuals, we cannot model the classification of individual volumes as Bernoulli trials. This is because the partitioning regime ensures that every volume is used once and only once as test data in each random partition set. Since the labels remain unchanged, there is in fact no randomness in terms of the test data, i.e.,

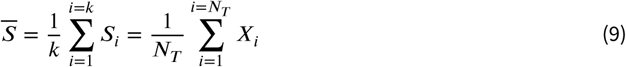

No matter how the data is partitioned, the pairing of *X_i_* and its label remains unchanged. Therefore *S*̅ is i constant and

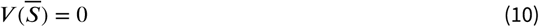

The problem here is that although the test data is identical over each complete partition set, the training data varies. That is, for *X_i_* in two partition sets, the corresponding training data differs. This difference creates variability in the classification outcome. For shuffled labels, this variability was irrelevant since classification outcomes were already assumed to be maximally independent. To account for the training set variability using true labels, we can reframe the problem as one where the test data is the reference, and we model how the training data varies with random partitions. Now the random partitions have substantial overlap so that only a small fraction are truly independent between partition sets. Fora given test data point *X_i_*, we can estimate the effective number of independent samples per fold, denoted as *n_f_*′. Following Equation 3,

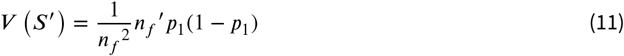

where *p*_1_ denotes the mean probability of success for that data set (volumes and labels combination). Using Equation 11 but otherwise following the same logic as the derivation of Equation 5, the variance of the distribution due to partition noise is estimated by:

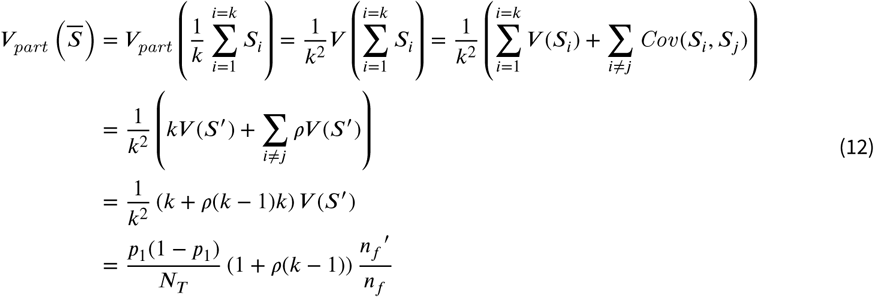

Now the factor *n_f_*′/*n_f_* is the fraction of data that is independent. Since the problem is reframed as one of variability in training data, the fraction is equivalently expressed as the fraction of training data that is independent, given a test data point *X_i_*. For large *k* and random partitioning, few of the remaining *n_f_* — 1 points in a fold with shared *X_i_* would be the same across partition sets. As a first order approximation, assume that all *n_f_* — 1 points are different, so that the fraction of distinct, and hence independent, data points in each training set is

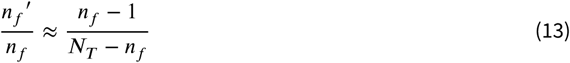

Substituting Equation 13 into Equation 12 we get:

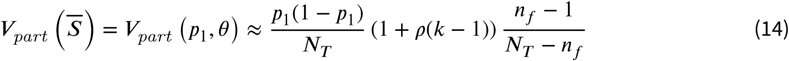

where the CV parameter *Θ* = (*N_T_*, *k*). For computational efficiency, we used a Gaussian approximation of the binomial model:

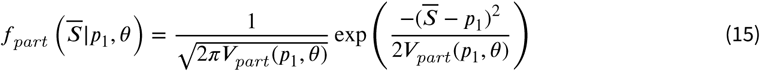

### Bayes Factor analysis

We defined a Bayes factor contrasting an alternative hypothesis with the null hypothesis:

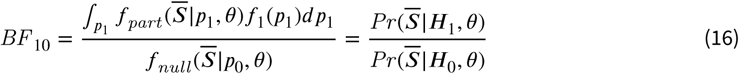

where the commonly used subscript _10_ denotes the alternative hypothesis is in the numerator and the null is in the denominator. Using the model for an individual’s true classification (unshuffled labels), we can compute the likelihood for the null hypothesis and the likelihood for the alternative averaged over a prior distribution *f*_1_. The typical prior distribution used is the most uninformative distribution that still converges for the Bayes factor calculation. For open intervals, that is usually the Cauchy distribution. In our case, classification rates cannot exceed 1, so the least-informative distribution is uniform between 0.5 (null) and 1, i.e.,

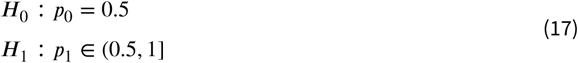

The uniform prior assumes that perfect classification success is equally likely *a priori* as just above chance. Although using the least informative prior potentially reduces unintended bias in the analysis, it also runs the risk of raising the threshold for finding evidence for the alternative, thereby seemingly favour the null. To test this possibility, two other prior distributions were also used for the alternative hypothesis, namely, a linear and quadratic distribution both maximal at *p* = 0.5 and decreasing to zero at *p* = 1. These distributions weight any alternative hypothesis *p* near 1 as less likely than the uniform prior.

For 0.5 < *p*_1_ ≤ 1, the three prior probability density functions of *p*_1_ used were

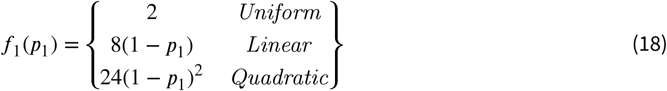

The density functions of Equation 18 were substituted one at a time into Equation 16, and combined with Equation 8 and Equation 15 to estimate the Bayes factor Equation 16. Note that for computing Bayes factor for location classification, *Θ* = (120,10), and for task classification, *Θ* = (240,10).

Assuming that *a priori*, the null hypothesis and weighted alternative hypothesis are equally likely, i.e., *Pr* (*H*_1_) = *Pr* (*H*_0_), then the Bayes factor is

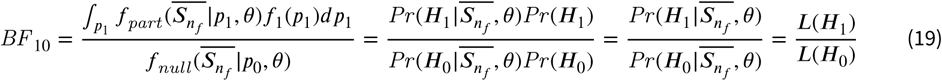

which is the relative likelihood of the alternative hypothesis to the null hypothesis, given the data and CV parameters. Consequently a large BF means more evidence for ***H***_1_, and a small BF means more evidence for ***H***_0_, as defined by *f_part_*, *f_1_* and *f_null_*.

## Results

### Behavioural performance

We set a stringent *a priori* performance criterion of 90% accuracy, to ensure that the participants were oriented during the task. This was necessary to minimize the possibility that failure to decode location from fMRI data could be due to poorly oriented participants.

The 18 participants included in the fMRI analysis had an average performance accuracy of 98.25 ± 0.56% (mean ± SEM). Remarkably, 50% ofthe participants did not commit a single error in 120 trials. Furthermore, the accuracies for target location 1 (mean ± SEM, 97.5 ± 0.8%) and target location 2 (98.9 ± 0.4%) were indistinguishable (p = 0.07, w_9_ = 38, Wilcoxon signed rank test), as were response times (mean ± SEM, 0.72 ± 0.03 s for target location 1, 0.75 ± 0.03 s for target location 2; p = 0.09, w_18_ = 124, z = 1.7, Wilcoxon signed rank test).

### Multivariate ROI analysis

Despite behavioural data demonstrating that participants were spatially oriented during the task, the multivoxel classifier could not predict location based on right hippocampal fMRI data. Figure 2a depicts a typical participant’s results for the classification of location, using our default method (i.e. LMGS detrending, 3 mm Gaussian smoothing, LSVM). As expected, the accuracy following random label-shuffles was distributed around the theoretical chance level of 0.5, since the shuffle process removes true location information. If multivoxel patterns were predictive of location in the virtual arena, then accuracies ofthe unshuffled data sets should be at or beyond the positive extreme ofthe shuffled distribution. Instead, unshuffled distributions were centred within the shuffled null distribution in all participants, arguing against the presence of location information at the voxel level. Notably, the variability in the unshuffled distribution can only be due to random partitioning itself since the set of unshuffled labels is unique. Thus if only a single partition is used, which is standard practice currently, it is unclear to which part of the partition distribution it might correspond (Figure 3a, red distribution). Therefore, to account for partitioning noise, statistical inferencing using cross-validation methods should be based on a sample of random partitions, or at least incorporate an estimate of partition noise variance. Using the default method, the partition noise variance in our data was 24 ± 2% (mean ± SD, n = 18) ofthe corresponding null distribution variance. For normally distributed independent random variables, if the true null variance is 24% larger than assumed, there would be 7.8% false positives at p < 0.05, and 2.1% at p < 0.01 (2-tailed false positive 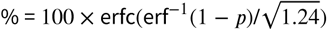), potentially inflating false positive conclusions by 1.5- to 2-fold.

**Figure 2.**
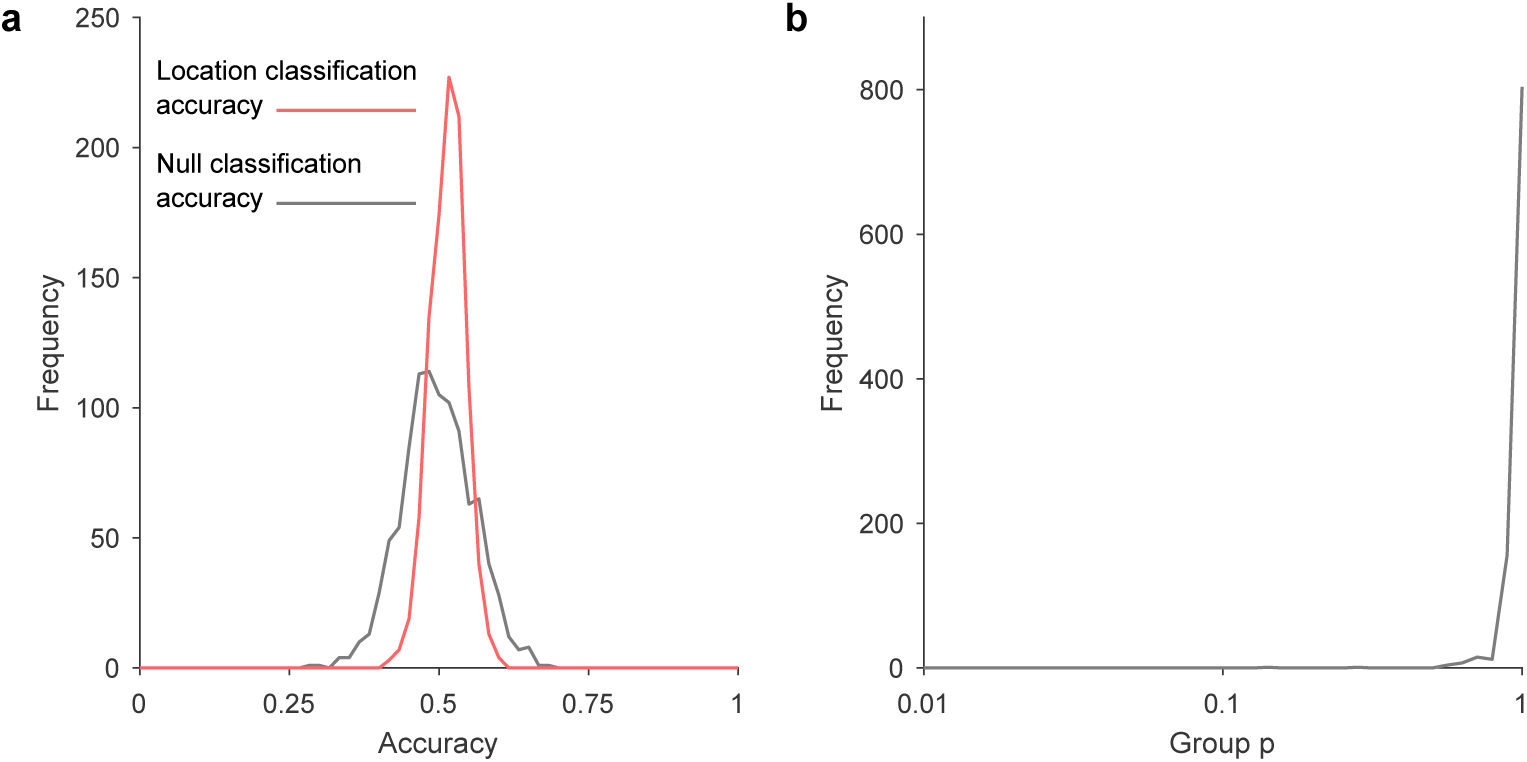
Results from right hippocampus for location classification. **a.** Atypical individual participant’s distribution of classification accuracies (10-fold stratified cross-validation results) for location in the virtual arena, over 1,000 random label-shuffles (black) and 1,000 random partitions of true labels (red). **b.** Population inference results for location classification following Allefeld et al (2016) show no evidence of a place code (18 participants, one p-value computed for each ofthe 1,000 random partitions).

**Figure 3.**
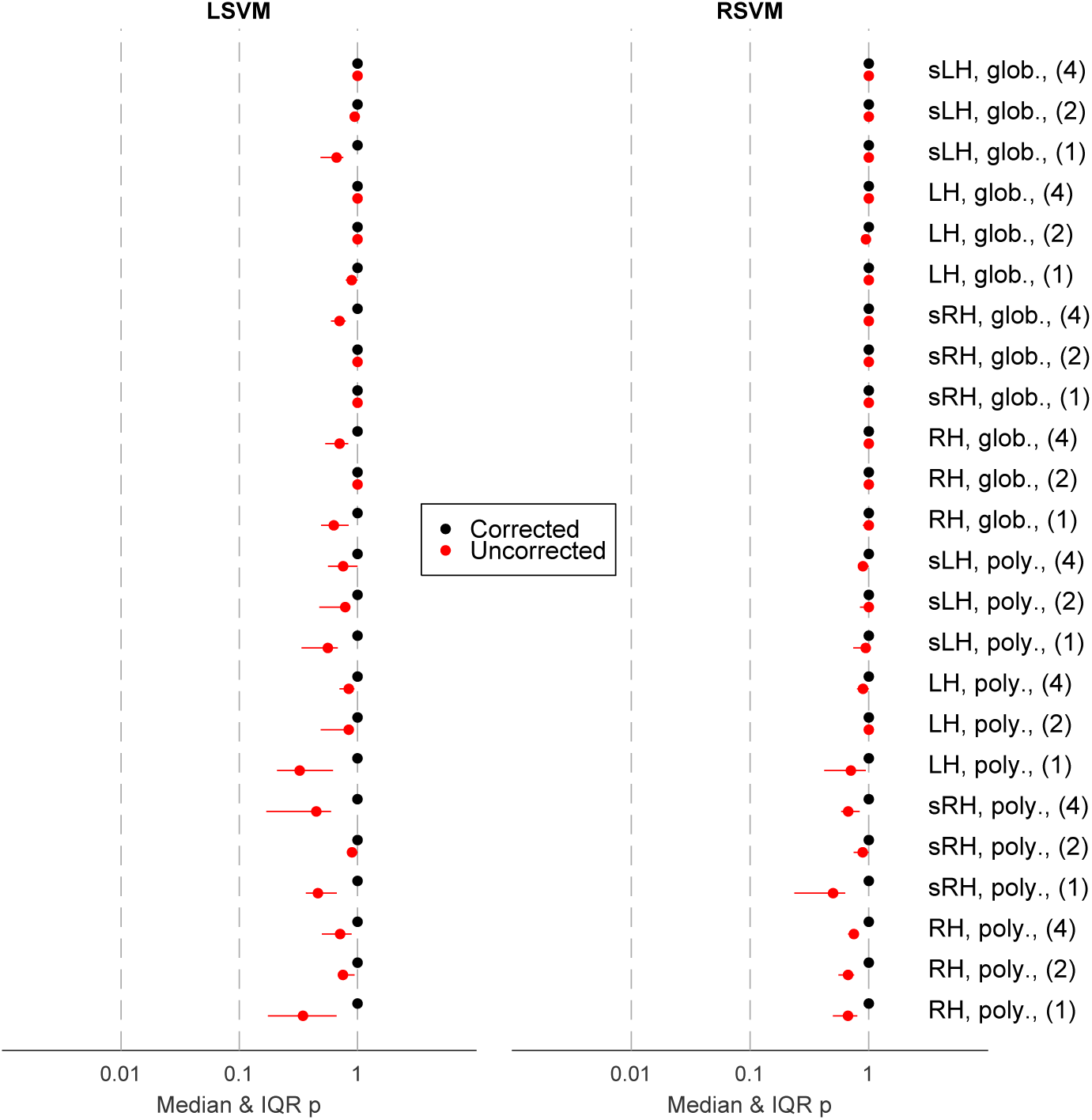
Overview of group significance results for different analysis approaches for the location classification following Allefeld et al. (2016), showing median as well as interquartile range. Abbreviations: glob. = Linear Model of the Global Signal detrending, H = hippocampus, L= left, R = right, LSVM = linear support vector machine, poly. = polynomial detrending (2^nd^ order), RSVM = support vector machine with radial basis function (Gaussian) kernel, s = smoothed (Gaussian kernel, radius = 3 mm). Numerals (i.e. 1,2, and 4) indicate number of consecutive images used for classification analysis.

For completeness, we submitted individual classification results from the 18 participants to a group analysis according to Allefeld et al. (2016). The prevalence null hypothesis states that the proportion of participants in the population having an above-chance location classification is zero. Figure 2b shows the group results for our default analysis where the group p > 0.1 for all random partitions, consistent with the null hypothesis that there is zero prevalence of location information in the population. Importantly, there was no evidence here that the conclusion may be affected by the instance of random partition of data used for cross-validation.

### Extended analysis of negative results

To investigate whether negative results could be due to our choice of preprocessing method, classifier, brain region or fMRI images (i.e. time period) we conducted several additional analyses to verify their validity. Figure 3 shows results for location classification across 24 different analysis approaches, including an alternative preprocessing method (second order runwise polynomial detrending), varying the number of consecutive images used for analysis, including left hippocampus, and including RSVM in addition to LSVM. Using LSVM, the median corrected group level p-value for the location classification under the prevalence null hypothesis exceeded 0.05 in all cases (Figure 3, left). In fact, even the lower limit ofthe 95% confidence interval ofthe p-value (arising from partition noise) exceeded 0.05. The same was true using RSVM (Figure 3, right). Our results also discount the possibility of a very weak but genuine voxel code that is by some means lost through the correction for multiple comparisons, since the median uncorrected p-value was never close to 0.05 (all p > 0.3). Therefore, no evidence for a classifiable voxel code for location was found, despite >98% mean behavioural orientation accuracy. Notably, there was no evidence that any particular choice of preprocessing method, classifier, ROI or timing made a significant improvement to location classification accuracy.

### Multivariate searchlight analysis

One possibility for a negative result may have been the “curse of dimensionality” because the data dimensionality (e.g. 3053 voxels in right hippocampus) is substantially higher than the number of data points available for classification (e.g. 60 visits to each location per participant). In fact, for both RSVM and LSVM, we found less than 1 classification error out of 120 when no data was withheld during training (averaged over participants, ROIs and preprocessing methods), showing that the problem was indeed of generalization to untrained data, rather than the separability of training data per se.

By restricting each classification problem to a small subregion of the ROI, searchlight analysis substantially reduces the data dimensionality, and has the potential to partially mitigate the dimensionality problem. Following Hassabis et al. (2009), we applied LSVM to spherical searchlights centred on each voxel in right and left hippocampus, and right and left parahippocampal gyrus (see *Methods* for details). This analysis produced 100 (cross-validation) accuracy values for each voxel of each ROI of each participant, using shuffled labels. Additionally, we produced an equivalent set of results from 100 random partitions of unshuffled data (for each voxel of each ROI of each participant).

Next we looked for evidence of a place code in any individual participants’ results using a nonparametric permutation analysis method (Nichols and Holmes, 2001). This approach avoids the need to make *a priori* assumptions about the data (which is implicit if statistical parametric maps are used). Beginning with the searchlight classification accuracy results, over each ROI, the maximum classification accuracy was found for each shuffled data set, and for each random partition ofthe unshuffled data set. We then found the number of random partitions (out of 100) for which the maximum statistic ofthe unshuffled searchlight results exceeded the 95% threshold ofthe shuffled searchlight results. If there is no signal, approximately five partitions should exceed the 95% threshold by chance. Across all ROIs, the mean number of partitions above the 95% threshold did not exceed 5/100 (mean ± SEM /100, RH = 3.2 ± 0.7, LH = 2.5 ± 0.8, RPH = 3.7 ± 0.7, LPH = 4.1 ± 1.1), showing no evidence of above-chance classification for location. We then asked whether it was possible that there could be a weak place signal which for some reason did not reach the arbitrary threshold of 95% ofthe shuffled data’s maximum statistic. We tested this possibility by counting the number of shuffled maximum statistics that each random partition’s unshuffled maximum statistic exceeded. The presence of a positive bias (>50%) may still suggest a weak but genuine place signal. Instead, no positive bias was found in any ROI (mean ± SEM, RH = 45 ± 3%, LH = 41 ± 3%, RPH = 44 ± 3%, LPH = 44 ± 3%).

In addition to the individual analysis, we also performed a group permutation test following Allefeld et al. (2016). Permutation-based information prevalence inference using the minimum statistic was used to determine if there is statistical evidence for a location code in the population (see Table 1). We started with the same searchlight classification accuracy results as above. In contrast to individual analysis, the minimum statistic was first found for all searchlights across participants, in each ROI. We used 10,000 second-level permutations, each of which was a random sample of one shuffled data set from each participant (one permutation was the unshuffled data). The minimum accuracy was found across participants, for each searchlight of each permutation.

**Table 1.** Group permutation test results showing the number of voxels for which p < 0.05 in each ROI, averaged across 18 participants.

| ROI | Mean ± SD no. voxels<br>$p < 0.05$, uncorrected | Mean ± SD no. voxels with<br>$p < 0.05$, corrected | Total no. voxels (common<br>to all participants) |
| --- | --- | --- | --- |
| RH | 74 ± 14 | 0.01 ± 0.10 | 2533 |
| LH | 91 ± 17 | 0.01 ± 0.10 | 2505 |
| RPH | 73 ± 14 | 0.01 ± 0.10 | 2157 |
| LPH | 59 ± 14 | 0.02 ± 0.14 | 1720 |

For each voxel, the uncorrected p-value was the fraction of permutation values of the minimum accuracy that was larger than or equal to the unshuffled data. Hence if the unshuffled accuracy is very high, very few of the permutation values will exceed it (low p-value). Since one permutation was the unshuffled data, the minimum p-value was 10^−4^. Even without correction for multiple comparisons, we found p < 0.05 in fewer than 4% of voxels in each ROI.

To correct for multiple comparisons (multiple searchlights), the maximum statistic (across searchlights) of the minimum accuracy (across participants) was computed. The p-value of the spatially extended global null hypothesis was the fraction of permutations in which the maximum statistic was larger than or equal to the unshuffled data. Across all random partitions, on average < 1 voxel reached p < 0.05 in each ROI (Table 1). Taken together, both uncorrected and corrected group results argue against the presence of location information in the searchlight accuracy values.

There remain a number of possible reasons that a place signal may not have been detected using the ROI-based and searchlight based multivariate classification methods described. One possibility is that the signal-to-noise ratio is too small to allow signal detection given the size of the training sets used for the classifier, or the number of participants tested in the case of group results. However, a number of studies have been reported that seemingly showed a voxel-level place signal using even fewer training points per participant, and fewer participants overall (Hassabis et al., 2009; Kim et al., 2017; Rodriguez, 2010; Sulpizio et al., 2014). Another possibility is that the analysis itself may be suboptimal for detecting this type of signal. To test this second possibility, we applied the difference-of-proportions analysis of Hassabis et al. (2009) to our searchlight accuracy values.

First, 10-fold stratified cross-validation results were pooled across all voxels in each ROI over 100 replications where location labels were randomly shuffled. This represents a null distribution of searchlight-based classification accuracy values, devoid of location information. For each ROI, the number of unshuffled voxels whose classification accuracy exceeded the 95^th^ percentile of the pooled distribution was found (Hassabis et al., 2009). The difference in the proportions of suprathreshold voxels was computed between all ROI pairs. According to Hassabis et al. (2009), finding a single proportion from each ROI avoids the problem of multiple comparisons across many searchlights within each ROI. We therefore replicated the analysis of Hassabis et al. (2009) immediately below, but show later that the implicit assumption of independence between searchlights is flawed.

Surprisingly, approximately half of all ROI contrasts resulted in p < 0.05 (Table 2). This suggests that the proportions of suprathreshold voxels differed between ROIs more than might be expected by chance. If the analysis is valid, this result may well imply that a multivariate voxel pattern exists in some (yet unexplained) location-and ROI-dependent manner. However, by virtue of including 100 random partitions, we could apply the same method to contrast two instances of the same ROI (diagonal cells of top-right section of Table 2). Clearly, a valid test should not detect a significant difference between the suprathreshold proportions arising from two random partitions of identical unshuffled data from the same ROI. Yet even for the same ROI, about half of all contrasts had p< 0.05. This suggests the false positive rate is at least an order of magnitude higher than it ought to be. On more careful inspection of the statistical methods used by Hassabis and colleagues, it becomes evident that the major reason is an underestimation of the test statistic’s standard error.

**Table 2.**
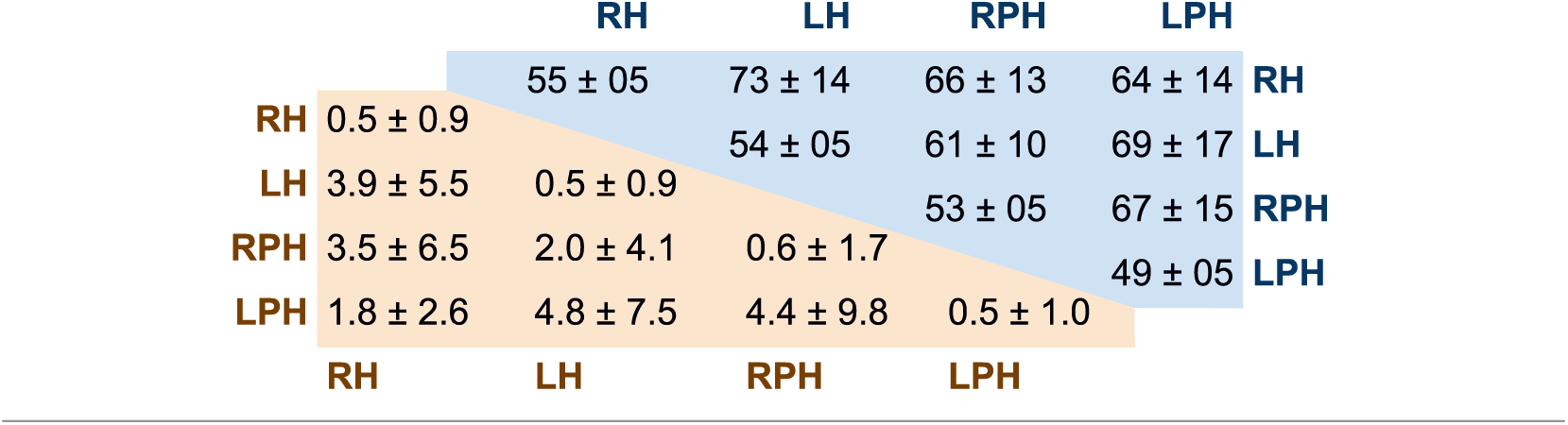
Percentage of ROI contrasts with p < 0.05 (top-right) difference-of-proportions method, 10,000 contrast pairs per participant, 18 participants (bottom-left) using shuffled data to estimate standard error of suprathreshold proportions, 10,000 contrast pairs per participant, 18 participants.

### Evaluation of analysis used in Hassabis et al. (2009)

Hassabis et al. (2009) compared the proportions of suprathreshold voxels identified through their standard searchlight analysis, from different ROI pairs. They then employed a commonly used formula (Daniel and Terrell, 1994) to estimate the standard error of the difference between two proportions, namely,

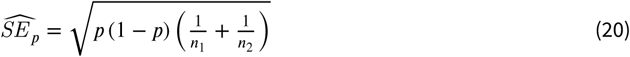

where the pooled proportion p is estimated by

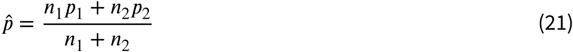

where n_1_ and n_2_ are the numbers of voxels in the two regions being contrasted, and p_1_ and p_2_ are the proportions of suprathreshold voxels in those regions. Using the estimated standard error from Equation 20, a Z-statistic was found which was then used to estimate the probability of a Type I error.

Using the estimated standard error from Equation 20 is incorrect here because the implicit assumption that independent Bernoulli-type outcomes contributed to the proportions being compared is violated. The proportion of suprathreshold voxels depends on the number of searchlights whose classification rates exceeded some threshold. However, each searchlight consists of a subpopulation of voxels, with substantial overlap with neighbouring searchlights. Therefore, the information in searchlights cannot be considered as independent. Indeed if one searchlight shows high classification accuracy, neighbouring searchlights that consist of many of the same voxels are also likely to show similar classification rates. In addition to the overlap of voxels between searchlights, neighbouring voxels themselves are known to show correlated activity due to physiology (e.g., shared blood flow) and preprocessing (e.g. low-pass filtering) (Poldrack et al., 2011). Empirically, we found a clear positive correlation between the classification accuracies of neighbouring voxels in right hippocampus (r = 0.72), right parahippocampal gyrus (r = 0.74), left hippocampus (r= 0.74), and left parahippocampal gyrus (r = 0.74). Neighbouring voxels were those centred no more than one voxel width away (i.e. maximum of eight neighbours) and within the same ROI mask. Correlations were computed between the mean accuracies of neighbouring voxels and the accuracies of the actual voxels themselves.

The assumption of independence between voxels therefore neglects the positive correlation between voxels, which leads to underestimation of the standard error of the difference in supra-threshold proportions. This in turn leads to underestimation of the probability of a Type I error. To test if the underestimation of the standard error of the difference-of-proportions was the major reason for the high percentage of ROI contrasts with p < 0.05 (Table 2), we re-estimated the standard error directly using the shuffled searchlight data. Using the same thresholding method as before, we computed 100 different supra-threshold proportions for each ROI (corresponding to all the shuffled data). Hence, for each ROI contrast, there were 100 difference-of-proportion values from shuffled data, used to estimate the mean and standard error of the null difference-of-proportions for that ROI contrast. For the same ROI pair (e.g. RH vs. RH), the standard error was estimated as the RMS of the other ROI pairs involving that ROI (e.g. RH vs. LH, RH vs. RPH, RH vs. LPH). As before, a Z-statistic was calculated, and a two-tailed p-value estimated using a normal approximation. Using this simple estimate of the standard error of the difference of suprathreshold proportions, the mean percentage of ROI contrasts with p < 0.05 dropped to less than 5% (Table 2, bottom-left). These results show that by using a more direct estimate of the standard error of difference-of-proportions, the percentage of contrasts with p < 0.05 is no more than expected by chance, arguing against an ROI-specific place code.

### Simulating faulty searchlight analysis using independent noise

It is unclear how much of the correlation of searchlight accuracies is a result of searchlight overlaps perse, and how much is a result of other factors such as shared blood flow or low-pass filtering which produces correlations in BOLD signal. It may be that overlaps between neighbouring searchlights contribute minimally to the underestimation of the standard error. If so, the problem should not exist if the underlying voxel data is truly independent. To investigate this possibility, we repeated the Hassabis analysis on pure noise. We generated 100 independent synthetic data sets by using Gaussian noise of the same mean, standard deviation and spatial distribution as voxels in our human fMRI ROIs, assuming statistical independence between all voxels. Analysis parameters were the same as for fMRI data. Note that the synthetic data sets were genuinely independent rather than merely using label shuffles as is the case for fMRI data. Since there was no true signal, we systematically excluded one data set at a time to simulate‘unshuffled’data (which should not be classifiable). By pooling the voxels from the remaining 99 data sets, we set the 95^th^ percentile threshold for classification accuracy as before. The number of voxels exceeding threshold in each of 100 ‘unshuffled’ data sets were used along with pooled proportions, and the standard error of pooled proportions to calculate Z-statistics. Using Gaussian approximation, we estimated 2-tailed p-values of the Z-statistics. For each ROI contrast, all 10,000 possible pairs of data sets were used (100 random partitions from each ROI).

If searchlight overlaps perse do not make a significant contribution to the correlation in searchlight accuracies, then there should be approximately 5% false positives (by setting p < 0.05) in the synthetic data. Instead, using the Hassabis method, there were more than 50% false positives in all ROI contrasts, including same-ROI contrasts (Figure 4 and Table 3), demonstrating that searchlight overlaps alone inflate false positive rates by an order of magnitude. Therefore, the searchlight method itself introduces enough correlation between otherwise independent voxels to violate the assumption of independence required to use uncorrected estimates ofthe difference-of-proportions. Taken together, our theoretical and experimental results demonstrate that the implicit assumption of independence in searchlight analyses by using uncorrected estimates of standard error of difference-of-proportions substantially increases false positives, and must be avoided.

**Figure 4.**
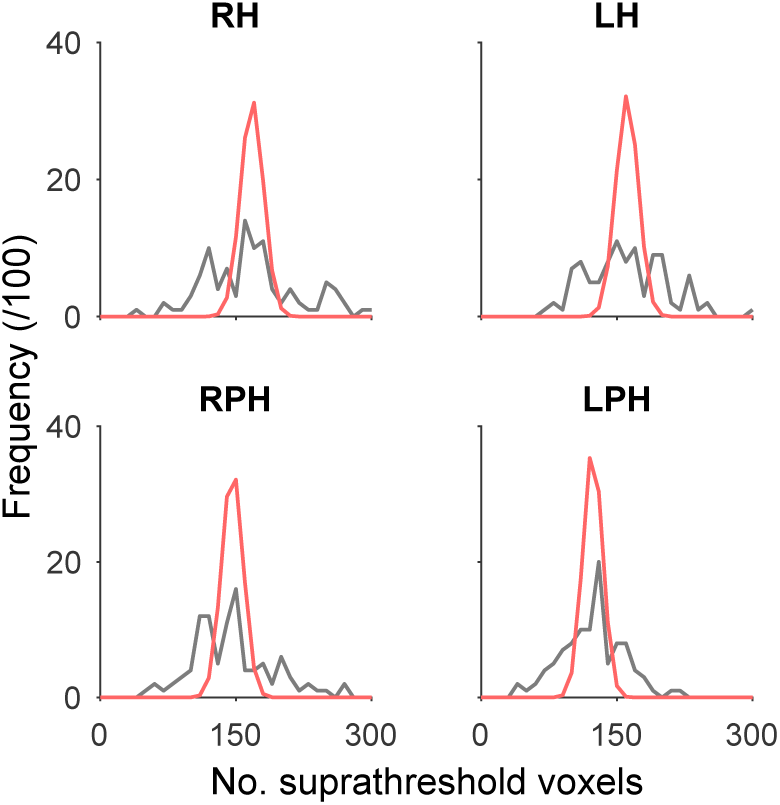
Frequency distribution of suprathreshold voxels in synthetic noise data sets corresponding to each individual ROI (black line, n = 100, see text for details). Using the same mean and assuming independent searchlight accuracies, a Gaussian approximation of the expected frequency of suprathreshold voxels (red line) shows substantial underestimation of the spread of suprathreshold voxel counts, causing an inflation of false positives, i.e. either higher or lower classification accuracies than expected by using the faulty null.

**Table 3.**
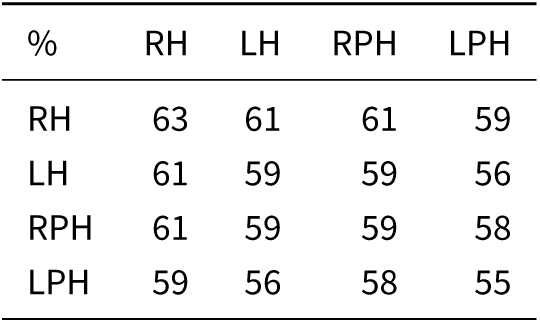
Percentage of ROI contrasts with p < 0.05 (pure noise example, difference-of-proportions method, 10,000 contrast pairs).

| % | RH | LH | RPH | LPH |
| --- | --- | --- | --- | --- |
| RH | 63 | 61 | 61 | 59 |
| LH | 61 | 59 | 59 | 56 |
| RPH | 61 | 59 | 59 | 58 |
| LPH | 59 | 56 | 58 | 55 |

### Positive control analyses

Since no evidence of a voxel-level place code could be found using a variety of approaches, we investigated the possibility that there was some unforeseen flaw in the image acquisition or analysis protocols. Using the same data, we determined whether two distinct phases in each trial, namely navigation vs. rest, could be classified (see *Methods*). Using our default method (i.e. LSVM, 3 mm smoothing, LMGS detrending) the two phases were clearly separable at a typical individual level (Figure 5a) and at the group level (Figure 5b). These analyses validate our image acquisition and data analysis protocols, and stand in contrast to our unclassifiable location results (Figure 2).

**Figure 5.**
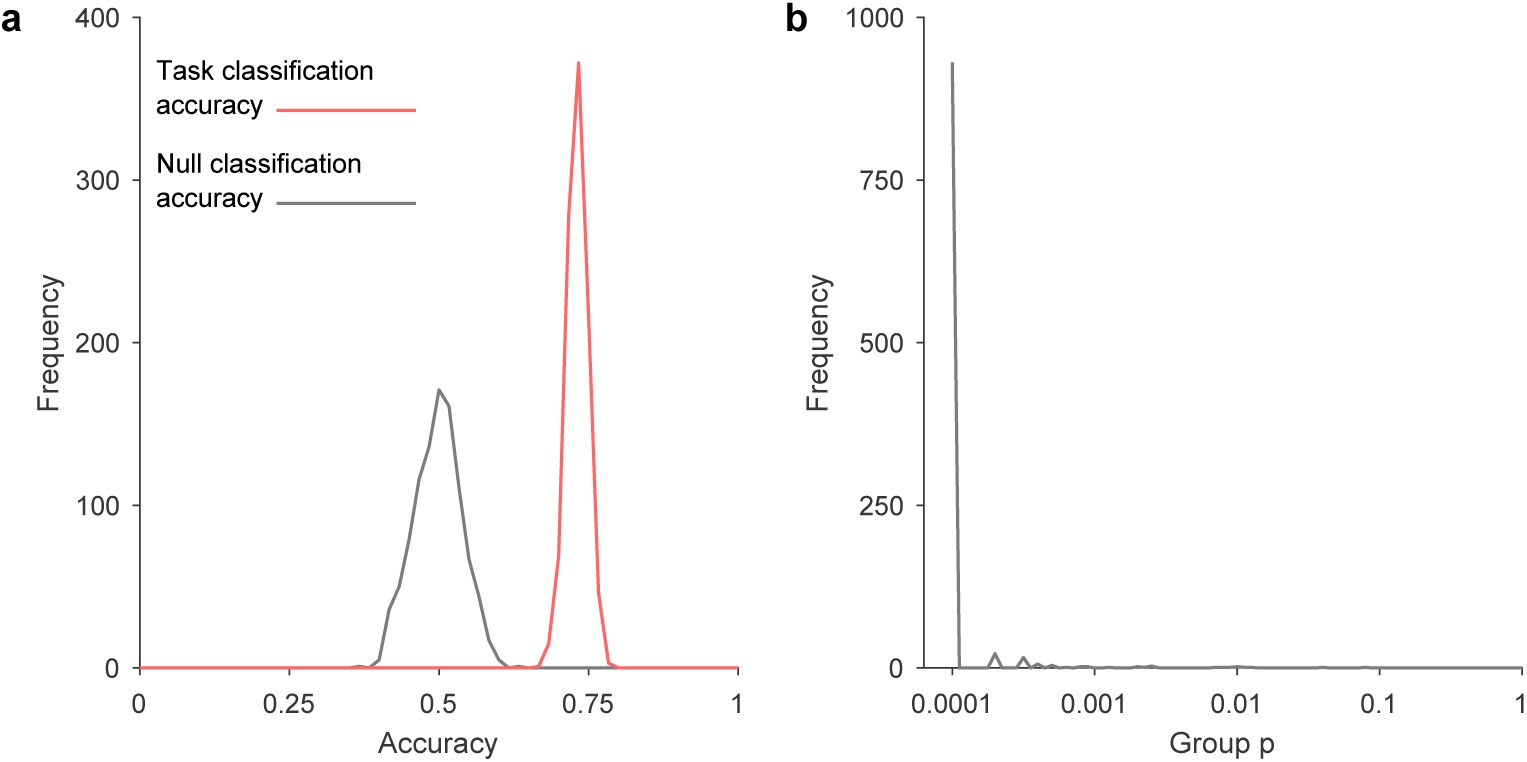
Results from right hippocampus for the control classification. **a.** Atypical individual participant’s distribution of classification accuracies (10-fold stratified cross-validation results) fortasktype (active vs. passive), over 1,000 random label-shuffles (black) and 1,000 random partitions (red) oftrue labels. **b.** Population inference results for control classification following Allefeld et al (2016) (18 participants, one p-value computed for each of the 1,000 random partitions).

Figure 6 shows results forthe positive control classification across 24 different analysis approaches. The median corrected group level p-valueforthe prevalence null hypothesis was less than 0.05 for all navigation vs. rest period classifications, across all ROIs, as well as smoothing and detrending methods, using LSVM (see Figure 6, left). The same was true of RSVM using polynomial detrending (see Figure 6, right). Note, however, that some 95% confidence intervals for the p-values included 0.05, showing that the choice of data partition can significantly affect classification generalization success. Nonetheless, for LSVM even the 97.5^th^ percentile p-value was below or close to 0.05 for both left and right hippocampus, using 2^nd^ order polynomial detrending. Thus at the group level, it is clear that voxel patterns are informative for rest vs. navigation periods of a task. Furthermore, we can exclude the possibility that only a small proportion of participants had classifiable voxel codes, which biased group results, since for all partitions where the null hypothesis was rejected, we can estimate the 95% confidence interval ofthe proportion of participants with a classifiable voxel code (Allefeld et al., 2016). Forthe smoothed right hippocampal data, LSVM resulted in null hypothesis rejection in 999/1000 random partitions. Of those, 0.62 to 1.00 of all participants are estimated to have a classifiable voxel code for rest vs. navigation (95% CI, median of partition shuffles). Taken together, these results suggest that hippocampal voxel patterns can be used to predict rest vs. navigation periods at above-chance level, in the majority of participants. Importantly, there is a clear difference between the classification performance for location 1 vs. location 2, and rest vs. navigation, using the same participants, experimental design, fMRI acquisition parameters, and analysis method.

**Figure 6.**
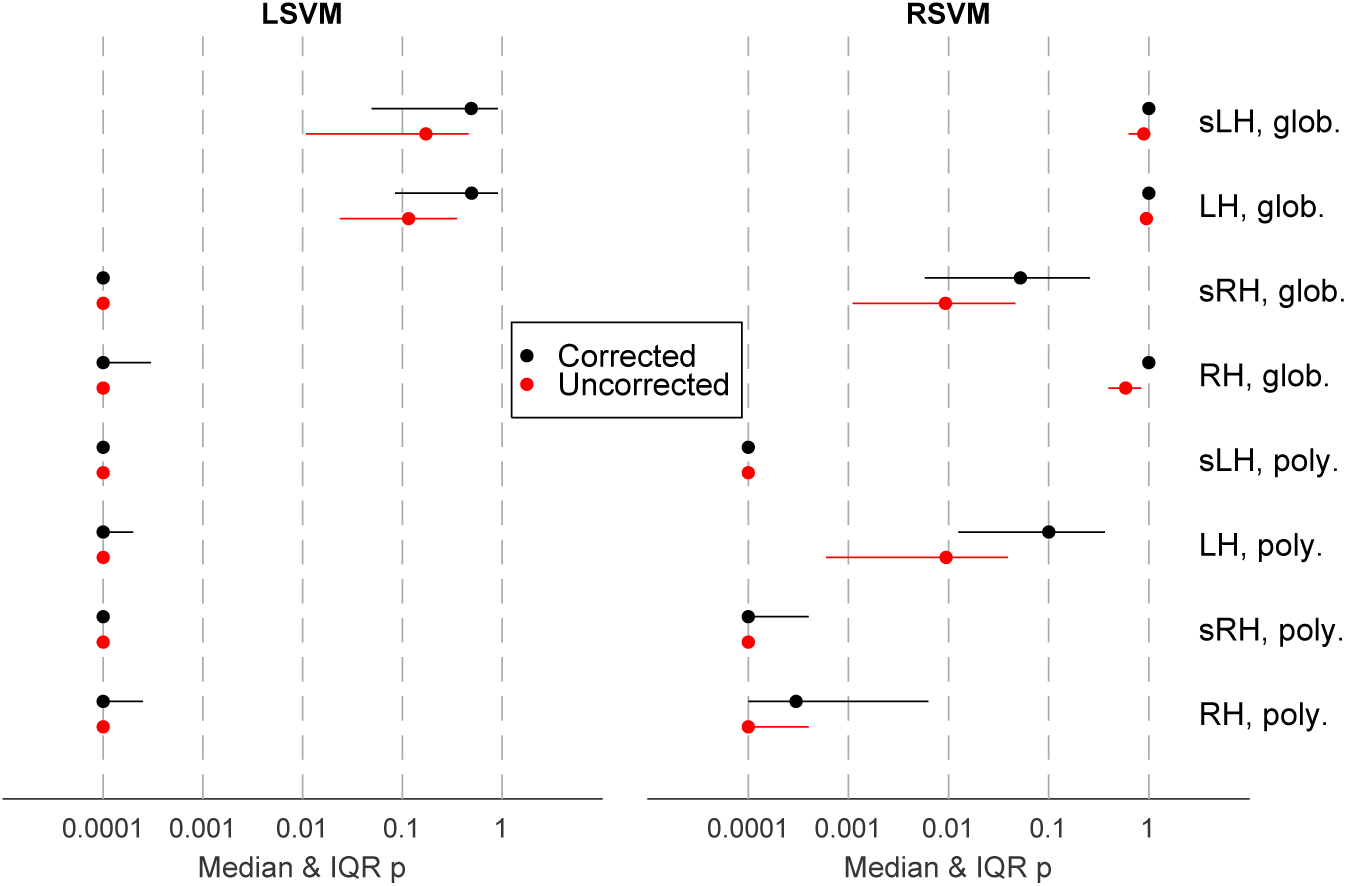
Overview of group significance results for different analysis approaches forthe control (i.e. task type) classification following Allefeld et al. (2016), showing median as well as interquartile range. Abbreviations: glob. = Linear Model ofthe Global Signal detrending, H = hippocampus, L = left, R = right, LSVM = linearsupport vector machine, poly. = polynomial detrending (2^nd^ order), RSVM = support vector machine with radial basis function (Gaussian) kernel, s = smoothed (Gaussian kernel, radius = 3 mm).

### Evidence for the null hypothesis

After careful analysis, we did not find any evidence to reject the null hypothesis that there is no voxel place code. However, finding no evidence to reject the null hypothesis is different to finding evidence to directly support it. Therefore, we considered whether the null hypothesis itself can be used to make testable predictions about the fMRI data. We used the default smoothed and globally detrended data from RH to test the predictions.

A straightforward prediction of the null hypothesis is that location labels do not matter and are effectively random when considering a population of participants. Thus for a sufficiently large sample size, the distribution of accuracies arising from true labels should be similar to the distribution due to shuffled labels. This was in fact the case for location classification (Figure 7a, red vs. black lines), where even distribution peaks arising from the discrete nature of scores were well matched. This directly supports the null hypothesis since true location labels were equivalent to shuffled labels, and were therefore uninformative. In contrast, if there is a genuine signal, then the two distributions should be distinct since the pooled distribution using true labels should no longer be equivalent to shuffled labels. This was in fact the case for task classification (Figure 7b, red vs. black lines), where the pooled distribution for true labels showed a higher mean and larger variance than for shuffled labels. These differences demonstrate that the true labels were not equivalent to shuffled labels, and therefore task information was present at the voxel level.

**Figure 7.**
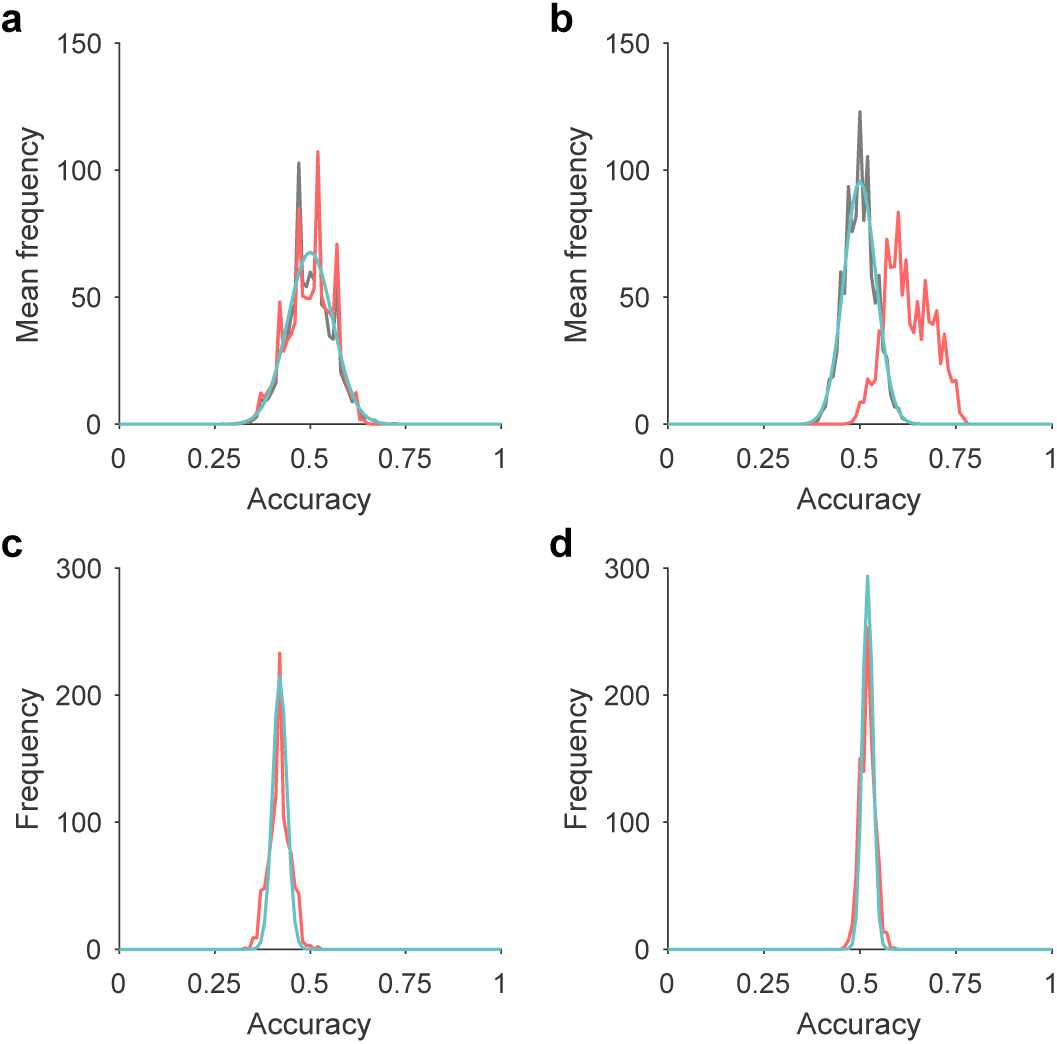
Comparison of noise model and LSVM accuracy distributions from RH. **a.** The frequency distribution of accuracy results is shown for location classification, averaged across all 18 participants, with shuffled (black) and true (red) location labels. A Gaussian approximation is shown (cyan) using a mean of 0.5 and variance estimated by a stochastic model assuming no label information. **b.** As per a butfortask classification. **c.** The frequency distribution of accuracy results is shown for location classification from a typical participant from a using true location labels (red). A Gaussian approximation is shown (cyan) using the mean of the individual’s sample, and variance estimated by a stochastic model assuming partition noise only. **d.** As per c butfortask classification.

Next we asked whether it is possible to derive an approximate form of the pooled distribution for location classification using true labels (Figure 7a, red line), using only the null hypothesis and experimental parameters. If so, this would show the null hypothesis is a sufficient model to account for the accuracy results, adding further evidence to support the null hypothesis for location classification. To do this we developed a simple stochastic binomial model of accuracy based on the null hypothesis (see Stochastic binomial model for shuffled labels). Our model was developed assuming statistical independence between data points which implies no label information. Hence our model should match data if there is no label information. Ourstochastic model provided a good match for location classification distribution with either true or shuffled labels (Figure 7a), suggesting that the null hypothesis provides a good quantitative account of location classification data. The stochastic model also predicts that the variance should be inversely related to the number of data points used for classification per participant. For task classification, there were twice as many volumes used for classification (two tasks per navigation sequence), and the pooled distribution for task classification using shuffled labels had a correspondingly smaller variance (Figure 7b).

To more directly contrast the evidence for the null versus alternative hypothesis, we computed Bayes factors for each participant’s accuracy results, using likelihoods estimated using models developed from the hypotheses. Therefore, in addition to the null model above, we needed a model of accuracy scores of individuals with true labels for the alternative hypothesis (that there is genuine information). Following simi-lararguments as above, we developed a simple stochastic binomial model of accuracy based on fixed labels and random partitions (see Stochastic binomial model for true labels, Figure 7c & Figure 7d). The model depended on the point accuracy score of classification as input, and predicted the corresponding accuracy density function. In this way, any prior distribution of accuracies can be used as the alternative hypothesis. To ensure that we did not inadvertently choose an alternative hypothesis which somehow biased outcomes, we tested three different prior distributions of accuracies reflecting varying prior beliefs about true accuracies (see Bayes Factor analysis).

There was a consistent pattern showing either no evidence (neutral) or evidence supporting (moderate, strong to extreme) of the null hypothesis for location classification (Table 4, location). In contrast, there was a consistent, but very different pattern showing either no evidence (neutral) or evidence supporting (moderate, strong to extreme) the alternative hypothesis for task classification (Table 4, task). Notably, the same pattern of results persisted across all three prior alternative hypotheses tested.

**Table 4.** Median Bayes factor (from 1,000 random partitions), out of a total of 18 participants, assumes shuffled labels variance for H_0_. Abbreviations: SH_0_ = Strong to extreme evidence for H_0_ (*BF* < 1/10), MH_0_ = Moderate evidence for H_0_ (1/10 ≤ *BF* < 1/3), N = Neutral (1/3 ≤ *BF*≤ 3), MH_1_ = Moderate evidence for H_1_ (3 < *BF* ≤ 10),SH_1_ = Strong to extreme evidence for H_1_ (10 < *BF*). BF category thresholds are based on Dienes (2014); Jarosz and Wiley (2014); Jeffreys (2000); Ly et al. (2016); Raftery (1995).

| Classification | Prior p distribution for $H_1$ | $SH_0$ | $MH_0$ | $N$ | $MH_1$ | $SH_1$ |
| --- | --- | --- | --- | --- | --- | --- |
| Location | Uniform | 8 | 7 | 3 | 0 | 0 |
|  | Linear | 5 | 4 | 9 | 0 | 0 |
|  | Quadratic | 5 | 3 | 10 | 0 | 0 |
| Task | Uniform | 0 | 1 | 6 | 2 | 9 |
|  | Linear | 0 | 0 | 6 | 2 | 10 |
|  | Quadratic | 0 | 0 | 3 | 5 | 10 |

Taken together, the convergence of distributional, model and Bayes factor results directly and consistently support the null hypothesis for location classification, and support the alternative hypothesis for task classification. These results complement the nonparametric population inference analyses to argue against any evidence for a voxel place code.

## Discussion

The goal ofthe present study was to reinvestigate whether human hippocampal place codes are detectable using fMRI. We employed a virtual environment that eliminated visual and path related confounds during the signal-decoding period to ensure that any positive finding would be indicative of a pure place code rather than a view code ora conjunctive view-trajectory code. We also employed a variety of signal processing and classification approaches, as well as a positive control analysis to evaluate carefully the possibility ofthe nonexistence of a purely spatial multivoxel place code.

Our experiment showed that, while participants were fully oriented during the navigation task, there was no statistical evidence for a place code, i.e. we could not reliably distinguish the two target locations using multivoxel-pattern classification algorithms. Additionally, we found robust and consistent evidence to directly support the null hypothesis for location classification data, using Bayes factor analysis and a model of SVM classification results derived from the hull hypothesis. These findings are in line with conclusions drawn from electrophysiological rodent data, which suggest that given the sparseness and distributed nature of place codes in the hippocampus, it would be implausible for them to be detectable using fMRI (O’Keefe et al., 1998; Redish and Ekstrom, 2012). Our findings are at odds with four prior imaging studies that reportedly have detected multivoxel place codes in the hippocampus (Hassabis et al., 2009; Kim et al., 2017; Rodriguez, 2010; Sulpizio et al., 2014). Since we employed a range of different image preprocessing and analysis approaches, it seems unlikely that our particular choice of analysis strategy could account for the discrepant results. Moreover, our control analysis showed that we were able to detect task-related changes in hippocampal activity, discounting the possibility that differences in image acquisition protocol or potentially image quality could be the reason prohibiting a positive finding.

In light of our results, it is important to carefully identify plausible reasons for the positive fMRI findings of published studies (Hassabis et al., 2009; Kim et al., 2017; Rodriguez, 2010; Sulpizio et al., 2014). We identified a number of shortcomings in the experimental tasks and analysis strategies of the four fMRI studies that could explain why each study seemingly detected a multivoxel place code in the hippocampus.

### Statistical issues

#### Invalid assumptions of statistical independence

Hassabis et al. (2009) made the implicit assumption of statistical independence between searchlight accuracies that is violated in fMRI data (see *Evaluation of analysis used in Hassabis et al. (2009)* for details). More detailed inspection of the suprathreshold counts from the original experiment (Hassabis, 2009, sec. 3.6.3 Sub-region dissociation) reveals that numerous suprathreshold proportions were in fact less than 5% despite using a 95^th^ percentile threshold. For example, for their pairwise location comparison for subject 2, the hippocampal suprathreshold count was 118/4032 searchlights (= 2.9%), the parahippocampal gyrus suprathreshold count was 70/3822 searchlights (= 1.8%), and the reported p-value was 0.002 for this contrast despite so few searchlights reaching the shuffled data’s threshold. Importantly, all p-values reported were replicable using the faulty method outlined earlier. Across 16 contrasts reported, 22/32 suprathreshold proportions were less than 5%. Therefore, these original results showed no evidence that location classification was possible in either ROI, and in hindsight should have raised alarms about the subsequent statistical methodology.

#### Paired t-test on accuracies

Rodriguez (2010) and Sulpizio et al. (2014) relied on a paired t-test for group analysis of decoding performance. As discussed in the *Methods*, when applied to classification accuracies, such a test will with high probability yield ‘significant’ results even though only a small minority of participants in the population shows above-chance classification (Allefeld et al., 2016).

#### Classifier confounds

Rodriguez (2010) included both the encoding and test phases of each trial in the dataset as independent trials. The classifier may have identified the general relatedness of the two phases being part of the same trial, rather than the spatial location per se. Many factors unrelated to location in the virtual arena could have contributed to two consecutive phases of a trial being similar, including simply being close in time.

Similarly, Sulpizio et al. (2014) included several identical images in the training and test sets (i.e. three instances per unique view were used for training the classifier and one for testing it in their leave-one-out cross validation procedure). This alone could lead to successful overall classification.

Finally, Kim et al. (2017) provided few details regarding the path structures used in the navigation task. It is only mentioned that pseudorandom trajectories were used and that 76% of all trials involved the inner eight (out of 64) locations used for the fMRI analysis. It is not clear from the description in which order the locations were visited. The nature of the trajectories could, however, have a significant effect on similarity of the fMRI signals associated with each location, either due to different levels of autocorrelation, or related to different levels of locational awareness that might be confounded with certain path characteristics. In short, without careful quantification of the path structure it is difficult to exclude the possibility that it might have contributed to the statistical discriminability of the fMRI signal associated with different locations.

### Experimental design issues

A true place code should be demonstrably selective for position in a mnemonic representation of space rather than particular visual cues. Unlike rodent place cells, however, earlier monkey work showed that primate hippocampal cells signal locations or objects being looked at, independently of current self-location (e.g. Robertson et al., 1998; Rolls, 1999; Rolls et al., 1997). These results show that mammalian hippocampal spatial codes are not necessarily place-specific, and in some cases may be intrinsically interwoven with visual inputs. Furthermore, electrophysiological recordings from the human hippocampus suggest that the majority of active neurons are not spatially-selective, but may instead respond to various types of visual stimuli (Kreiman et al., 2000). Unfortunately, all four studies that claim to provide evidence for a voxel place code (Hassabis et al., 2009; Kim et al., 2017; Rodriguez, 2010; Sulpizio et al., 2014) failed to remove significant visual confounds, which implies that even a legitimate voxel code in these experiments could be sensory-driven rather than be a true place code.

#### Visually distinct landmarks

Reliable and unique visual landmarks pose a particular problem. In the most obvious scenario, such a cue might be visible in a period used for classification, a possibility in the study by Rodriguez (2010) (depending on the field of view). Even in the case that the cue is not visible at the classification point, however, visual traces or visual memory could account for positive classification. Participants in the Rodriguez (2010) experiment took direct paths to the goals (time limited, active navigation task), therefore the egocentric view direction of the landmark during navigation varied systematically with the goal location. Similarly, the virtual environments used by Hassabis etal. (2009) consisted of visually distinct landmarks on or adjacent to all walls, visible en route to target locations. The virtual environment outlined by Kim et al. (2017) contained a salient local landmark (a green door, whose visibility depended on the direction of travel and visual obstructions). The authors stated that the door was “occasionally” visible, but failed to demonstrate that neither those times nor visual appearance of the door were correlated with impending arrival location. In all three of these cases, above-chance decoding could be due to differences in visual information during navigation rather than pure spatial location.

#### Visual panoramas

Unique landmarks are a specific case of the more general problem of unique panoramas: any unique classifiable code may be due to the particular combination of visual cues rather than the more abstract notion of place. Such a confound was present in the experiment by Sulpizio et al. (2014), which required that static visual scenes completely determine location and orientation. Similarly, Kim et al. (2017) compared parallel locations in their rectangular environment, which would undoubtedly provide different panoramas — independent of the allocentric direction — due to different wall distance configurations. This problem could have been avoided by keeping the lattice edges rotationally symmetric but comparing only diagonally opposite rather than parallel corners, as these are visually equivalent (2-fold rotational symmetry; Cheung, 2014; Cheung et al., 2008; Stürzl et al., 2008)).

#### Optic flow

In the study by Kim et al. (2017), there was a connection bias between the locations in the 3D environment employed (i.e. not every location was connected to every location, and connections were not always symmetric) that caused the optic flows to differ depending on which test location was immediately upcoming. Earlier animal studies have shown that the hippocampus is sensitive to visual aspects of linear and rotational motion (O’Mara et al., 1994), and that it receives information from the accessory optic system (Wylie et al., 1999), which is a visual pathway dedicated to the analysis of optic flow. Hence, a classifier may be able to detect differences in preceding ground optic flow, which in turn correlated with test location.

#### Correcting visual confounds

In an attempt to alleviate concerns regarding the visual cues, Kim et al. (2017) used a simple visual texture model (Renninger and Malik, 2004) that provided a single visual similarity value for each trial (i.e. images captured at every half a second during the five second journey period for each location were averaged and entered into the texture model). The authors showed that even if these visual similarity measures were included in the analysis, location could still be inferred from the anterior hippocampal voxel patterns, and took this as a confirmation forthe existence of a pure place code. It is, however, questionable whether the visual texture model was suited to account for the differences in visual scenes encountered during navigation. For example, having a long wall to the right and a short wall to the front defines a distinct location to having a long wall to the left and a short wall to the front. Yet visual textures and other low level visual features may be virtually identical. The only way to ensure that differences in visual information during navigation cannot affect voxel patterns is to eliminate them entirely from the task design.

### Conclusions

All existing studies which assert to have found evidence fora hippocampal place code using functional magnetic resonance imaging can be challenged based on either statistical or task-related concerns and provide no robust convincing evidence of a multivoxel place code in humans. Further evidence against the detectability of a hippocampal place code using functional magnetic resonance imaging comes from a published pilot study (n = 3) by Op de Beeck et al. (2013) which employed a virtual navigation paradigm with the aim of decoding location information from fMRI activation patterns, but also found no statistical evidence for a place code in the hippocampus. They were, however, able to statistically infer spatial location from voxel patterns in the visual cortex, giving further weight to our concerns regarding visual confounds in the aforementioned studies. Moreover, a number of recent studies have shown that patients with hippocampal damage have difficulties in complex visual discrimination task, suggesting a role ofthe hippocampus in visual perception (Hartley et al., 2007; Lee et al., 2005a,b, 2006,2007). In contrast, activity of bona fide place cells identified in rodents has been shown repeatedly to be view-independent and persists even without visual information (Quirk et al., 1990; Rochefort et al., 2011; Save et al., 1998, 2000). Hippocampal place cells of bats have also been shown to persist without visual input (Ulanovsky and Moss, 2007). Similarly, pure place cells identified in the hippocampus of epilepsy patients were also view-independent (Ekstrom et al., 2003). In line with place cell properties common to phylogenetically diverse mammalian species, claiming the existence of a multivoxel place code necessitates exclusion of direct visual contributions to activity differences.

In summary, we have conducted a detailed assessment ofthe claim that place codes are detectable using fMRI in human hippocampus. Our combined experimental and theoretical results provide rigorous and consistent evidence against this claim. Additionally, we identified several serious shortcomings in published imaging studies claiming evidence in favour of a hippocampal multivoxel place code. We also note that electrophysiological data suggest that hippocampal place codes are both sparse and anatomically distributed, so that imaging techniques such as fMRI should not, at least at present, be capable of detecting location-specific place cell activity. Taking all evidence in combination, claims of the existence of a purely spatial voxel code of location should therefore be treated with appropriate scepticism. We assert that any future imaging study claiming evidence in favour of a multivoxel place code should rigorously eliminate potential confounds due to visual features, path trajectories and semantic associations that could lead to decodable differences between spatial locations. In addition, it will be crucial to employ appropriate and robust statistical tools to avoid false positives that are a particular concern for high dimensional data.

